# JEPA-SORT: World-Model Embeddings Improve Deep-Sea Multi-Object Tracking on Visually Homogeneous Fauna

**DOI:** 10.64898/2026.10.07.757484

**Authors:** Amory Tillinghast-Raby, Matt Kravetz, Jeriyla Kamau-Weng, Alexis Weinnig, Yara Rodrigues, Filippo Varini, Vincent Pieribone

## Abstract

Protecting deep-sea sessile communities as Marine Protected Areas (MPAs) requires accurate estimates of their spatial distribution and abundance. Existing computer-vision detection methods require careful calibration to avoid counting errors; multi-object tracking (MOT) theoretically offers a more robust alternative. In practice, however, poor lighting and turbidity limit the efficacy of purely kinematic MOT algorithms in deep-sea environments. At the same time, the intra-species visual homogeneity typical of deep-sea scenes means traditional per-frame appearance features are often too weak to maintain stable identities in deep-association MOT algorithms. In this paper we introduce JEPA-SORT, an MOT algorithm that injects V-JEPA 2.1 embeddings into detection-to-track association, to show that video world-model embeddings - which capture not only physical appearance but spatiotemporal trajectory across viewpoints - can help maintain stable identities on visually homogeneous specimens. We provide a benchmark of twelve ROV transects surveying deep-sea sessile communities and evaluate JEPA-SORT against standard MOT algorithms across a range of simulated optical environments. We also propose the Biodensity Utility Metric (BDU), which measures tracker performance in the context of downstream ecological modeling. JEPA-SORT outperforms all benchmarked trackers on IDF1, MOTA, and BDU and is competitive on specimen-count accuracy. Ablations indicate its advantage stems primarily from the discriminative power of world-model embeddings on visually homogeneous footage.

## 1. INTRODUCTION

The deep sea makes up 95% of the ocean by volume yet remains one of the least explored biomes on Earth (Danovaro et al., 2017). In this environment, sessile organisms such as cold-water corals and sponges build three-dimensional reef structures that support entire ecosystems by trapping particulate matter and providing habitat, feeding grounds, and nurseries for diverse species (Maier et al., 2023). As both individual species and ecosystem engineers, they form the foundation of broader biological assemblages and warrant protection through Marine Protected Areas (MPAs), which depend on robust frameworks for describing their composition, distribution, and abundance.

Advances in imaging, including remotely operated vehicles (ROVs) and autonomous underwater vehicles (AUVs), now enable in situ observation of benthic organisms and their ecological interactions (López-Garrido et al., 2020), while machine learning (particularly computer vision) allows these organisms to be detected and identified in footage at scale. Identification alone, however, cannot capture abundance and distribution, both critical for assessing MPA status. Abundance is commonly estimated by counting identifications in frames sampled at a fixed interval that prevents overlap between samples (Durden et al., 2015). This minimizes overcounting, which artificially inflates abundance estimates, but it requires careful calibration with constant ROV/AUV speed and field of view, limiting the ability to focus on specific targets. Fixed-interval sampling also discards much of the transect, costing opportunities to identify organisms that may be unidentifiable in specific individual frames.

Multi-object tracking (MOT) theoretically offers a more robust solution by maintaining detection identities across continuous footage. In practice, however, poor lighting, turbidity, and intra-species physical homogeneity make robust tracking in deep-sea footage difficult (Liu et al., 2022; de Langis and Sattar, 2020; Wang et al., 2026). Furthermore, MOT metrics such as IDF1 and MOTA also fail to align cleanly with MOT algorithms’ utility for ecological assessment (Chan et al., 2026). And while MOT algorithms are acknowledged as suffering from these limitations, to our knowledge, no benchmark has compared MOT algorithms’ performance on deep-sea abundance estimation from ROV/AUV footage across varying environmental conditions.

This paper makes two key advancements. First we introduce the JEPA-SORT algorithm: a simple, online, and realtime tracker that injects embeddings from Meta’s V-JEPA 2.1 video world-model into detection-to-track association. Both an algorithmic and a conceptual contribution, the new JEPA-SORT algorithm is necessary to utilize embeddings from V-JEPA 2.1, which are calculated from 16-frame clips and therefore cannot be directly injected into an extant deep-association tracker like DeepSORT, that expects per-frame embeddings. Equally importantly, however, JEPA-SORT illustrates how video world-model embeddings, which encode both a speci-men’s physical appearance and spatiotemporal trajectory across frames, can substantially improve MOT performance in environments with significant detection dropout and high visual homogeneity. Second, we provide the Biodensity Utility Metric (BDU) which measures tracker performance in the context of downstream ecological modeling.

To facilitate this analysis, we provide an open-source benchmark of twelve ROV transects surveying deep-sea sessile communities (Instituto do Mar and OceanX, 2026). Compared to four standard MOT algorithms: SORT, OC-SORT, DeepSORT, and DeepOC-SORT, JEPA-SORT outperforms across standard MOT metrics such as IDF1 and MOTA (Heindl, 2017). It also leads on BDU and is competitive on raw specimen count accuracy. Ablations indicate that while the JEPA-SORT architecture provides a strong algorithmic framework, the algorithm’s competitive advantage is primarily due to the discriminative power of world-model embeddings on visually homogeneous footage.

## 2. MATERIALS AND METHODS

The following section is divided into five parts: the JEPA-SORT architecture, explanation of the oracle detector, the definition of BDU, a description of the provided benchmark, and the experimental design used to compare JEPA-SORT against standard MOT algorithms.

### 2.1. JEPA-SORT architecture

JEPA-SORT belongs to the simple, online, and realtime family of trackers, building on the architecture shared by SORT, OC-SORT, and their deep variants: detections are associated to tracks frame-by-frame, without knowledge of future states, by solving a linear assignment problem over the detection-to-track cost matrix (Bewley et al., 2016). Its key innovation is combining frame-level kinematic association with spatiotemporal appearance embeddings from a video world-model. Unlike the per-frame embeddings used in DeepSORT, spatiotemporal appearance embeddings encode both a specimen’s visual features and its trajectory across frames, making them especially well suited to deep-sea environments, where intra-species visual homogeneity is high.

#### 2.1.1. Overview

JEPA-SORT uses a standard, constant-velocity linear Kalman filter, *K*, with an eight-dimensional state (*u, v, γ, h, u*^′^*, v*^′^*, γ*^′^*, h*^′^), where (*u, v*) denotes the bounding-box center, *γ* the aspect ratio, and *h* the height (Wojke et al., 2017). Internally, each JEPA-SORT track is represented as a tuple (*k, h, g, a*), where *k* is the Kalman state, *h* is the track history, *g* is a gallery of associated spatiotemporal appearance embeddings, and *a* is the time since last update. Our implementation of JEPA-SORT utilizes Meta’s V-JEPA 2.1 model (Mur-Labadia et al., 2026). The core components of JEPA-SORT - track state management, association logic, and V-JEPA 2.1 feature-token pooling strategy, are detailed below. Table 1 summarizes the notation used throughout this section.

**Table 1:**
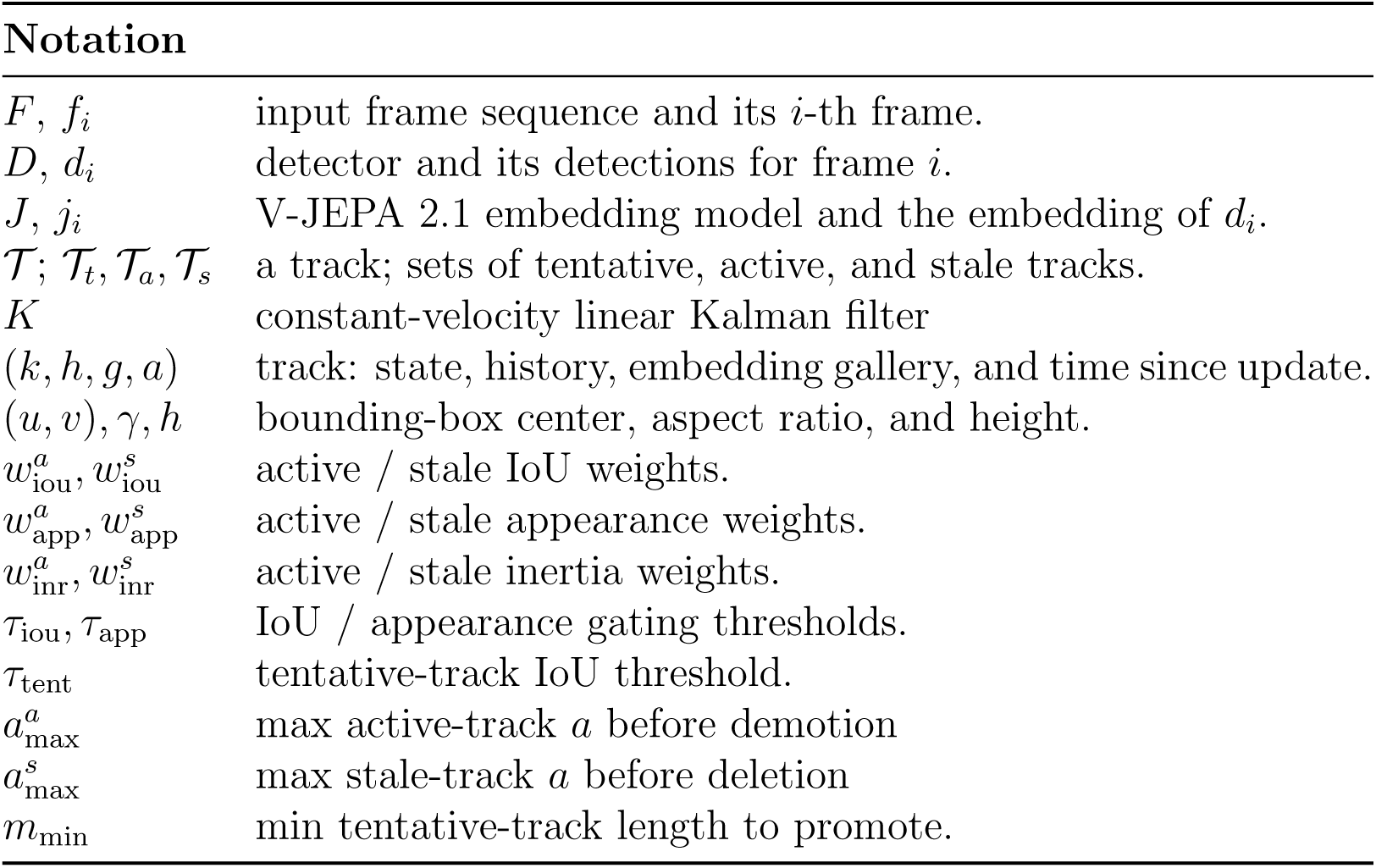
JEPA-SORT notation.

| **Notation** |  |
|---|---|
| F, f<sub>i</sub> | input frame sequence and its i-th frame. |
| D, d<sub>i</sub> | detector and its detections for frame i. |
| J, j<sub>i</sub> | V-JEPA 2.1 embedding model and the embedding of d<sub>i</sub>. |
| 𝒯; 𝒯<sub>t</sub>, 𝒯<sub>a</sub>, 𝒯<sub>s</sub> | a track; sets of tentative, active, and stale tracks. |
| K | constant-velocity linear Kalman filter |
| (k, h, g, a) | track: state, history, embedding gallery, and time since update. |
| (u, v), γ, h | bounding-box center, aspect ratio, and height. |
| w<sup>a</sup><sub>iou</sub>, w<sup>s</sup><sub>iou</sub> | active / stale IoU weights. |
| w<sup>a</sup><sub>app</sub>, w<sup>s</sup><sub>app</sub> | active / stale appearance weights. |
| w<sup>a</sup><sub>inr</sub>, w<sup>s</sup><sub>inr</sub> | active / stale inertia weights. |
| τ<sub>iou</sub>, τ<sub>app</sub> | IoU / appearance gating thresholds. |
| τ<sub>tent</sub> | tentative-track IoU threshold. |
| a<sup>a</sup><sub>max</sub> | max active-track a before demotion |
| a<sup>s</sup><sub>max</sub> | max stale-track a before deletion |
| m<sub>min</sub> | min tentative-track length to promote. |

#### 2.1.2. Track Management

To suppress false positives while remaining robust to occlusion, tracks move through three registries: tentative tracks (T*_t_*), active tracks (T*_a_*), and stale tracks (T*_s_*). Each registry has tunable hyperparameters that control the balance between kinematic and spatiotemporal appearance-based association; see Table 1 for details.

##### Algorithm 1

JEPA-SORT track management (high-level)

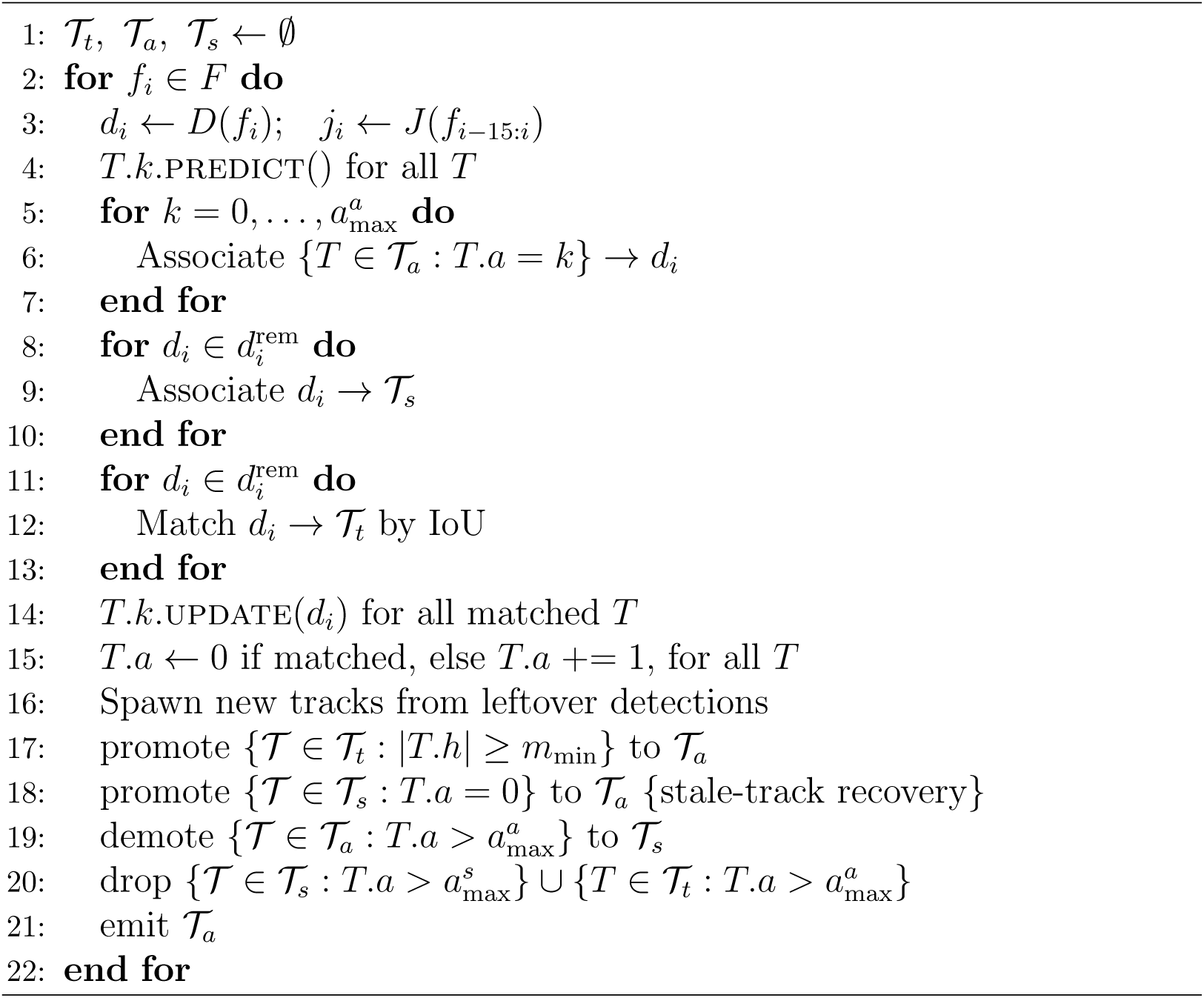

Track management proceeds as follows. At each timestep, the Kalman state of every extant track is advanced by prediction, and the incoming frame is processed by both the detector, *D*, and the V-JEPA 2.1 embedding model, *J*. The resulting detections are matched to tracks across the tentative, active, and stale registries using cascading association logic (see Section 2.1.3). Associated tracks are corrected by Kalman update, and each track’s time-since-update counter, *a*, is either reset or incremented. Tracks then transition between registries according to three rules:

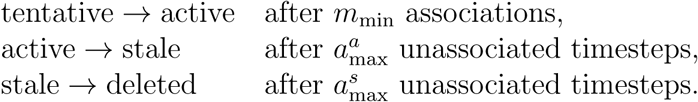

Stale-track recovery, re-promoting a stale track to the active registry after renewed association, is the primary mechanism by which JEPA-SORT handles prolonged occlusion and detection dropout.

##### Algorithm 2

Association

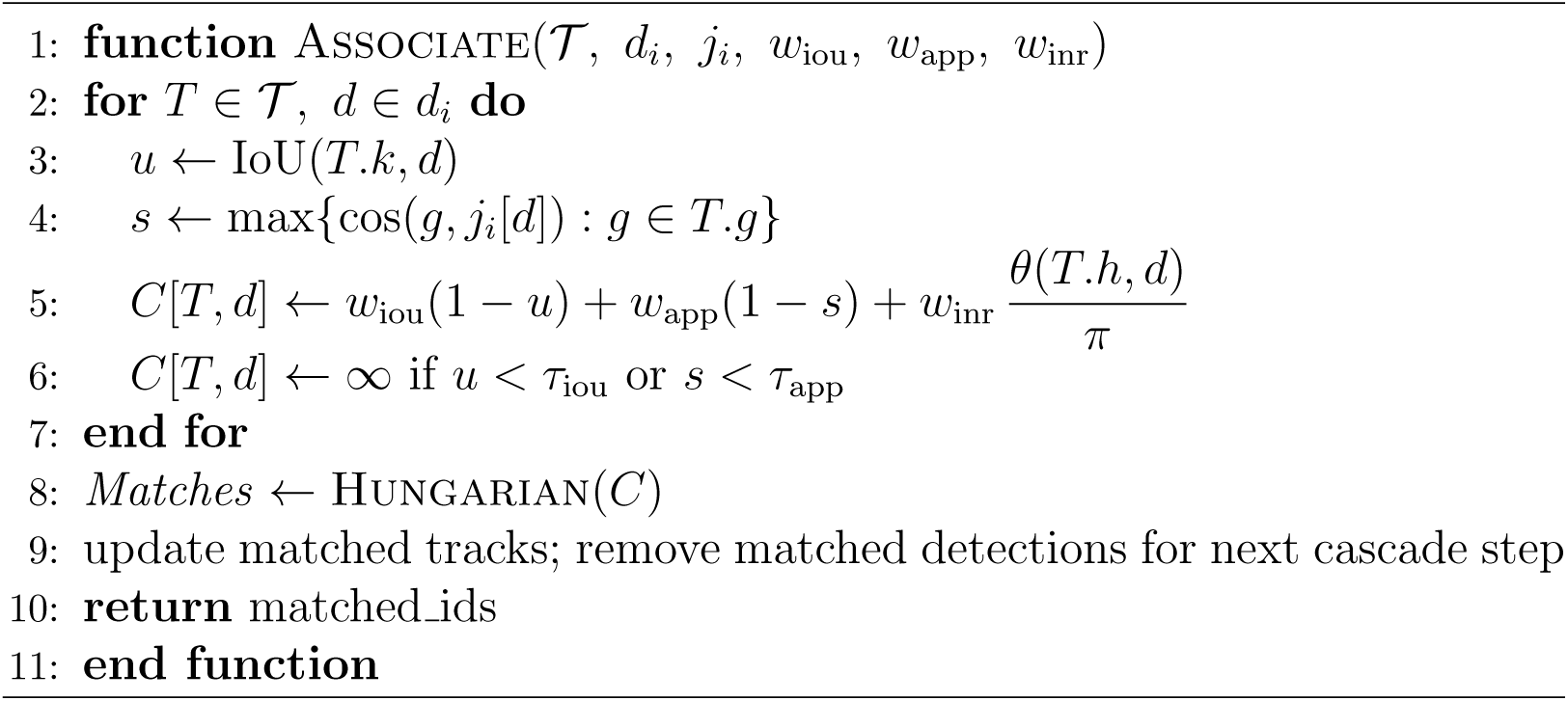

#### 2.1.3. Association

In JEPA-SORT, detection-to-track association is cascading (Wojke et al., 2017) and combines three terms: kinematics (detection-candidate IoU), spatiotemporal appearance (cosine similarity between a detection and candidate track’s embedding), and inertia (angle between a detection and candidate track). The inertia term, adopted from OC-SORT (Cao et al., 2023), penalizes associations that imply a large change in direction relative to the track’s previous state.

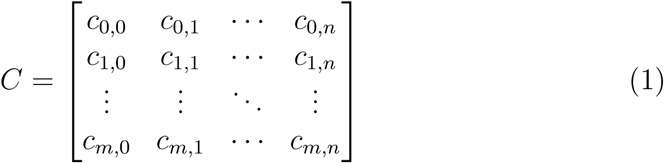

For active-track association, tracks are partitioned by time since last update, T *.a*. Association proceeds in descending order of recency, beginning with tracks where T *.a* = 0 and then continuing through T *.a* = 1, 2, For each subgroup, JEPA-SORT computes the cost matrix (1), and solves the assignment with Hungarian(*C*). For any candidate track-detection pair, the cost is set to infinity if either the kinematic term IoU(*T.k, d*) or appearance term max(cos(*T.g, j_i_*[*d*])) are less than their respective thresholds: *τ*_iou_ and *τ*_app_ (see Algorithm 2). Detections left unassigned remain eligible for the next active-track subgroup.

After active-track association, stale tracks are associated as a single group using the same procedure. Remaining detections are then matched to tentative tracks by greedy IoU association. Lastly, for any detection that is still unmatched, JEPA-SORT initializes a new track with its own unique ID.

#### 2.1.4. V-JEPA Feature Token Pooling

Unlike DeepSORT and other frame-based re-identification methods, JEPA-SORT uses embeddings that encode both appearance and spatiotemporal history over the 16 most recent transect frames. We are not aware of prior work applying V-JEPA embeddings to multi-object tracking.

To calculate the embeddings, each incoming frame is added to a sliding 16-frame window, which is passed into V-JEPA 2.1. Detection-level representations are then extracted by projecting detection bounding boxes onto the resulting spatiotemporal token grid, pooling the tokens within each detection over the most recent two-frame tubelet, and L2-normalizing the pooled vector.

At the start of each transect, before 16 frames are available, the clip is left-padded with 16 − *N* copies of the first frame, where *N* is the number of processed frames including the current frame.

### 2.2. Explanation of the Oracle Detector

To isolate tracker performance from detector performance, all trackers draw detections from a shared oracle rather than a specific trained model like YOLO. At each frame, the oracle returns ground-truth bounding boxes from MOT-formatted transect labels, with adjustable perturbations emulating underwater detector failure modes in varying environments: detection dropout, false-positive injection (with optional forward propagation to mimic persistent false detections from particulates or biofouling), and bounding-box area jitter.

### 2.3. Definition of BDU

Standard MOT metrics do not directly measure tracker performance in the context of ecological modeling, which requires both accurate specimen counts and consistent track identities (Chan et al., 2026). We therefore introduce the Biodensity Utility Metric (BDU), a composite of standard MOT metrics that jointly measure specimen-count accuracy and track-identity consistency while scaling performance based on the number of unique ground-truth organisms - a proxy for scene complexity.

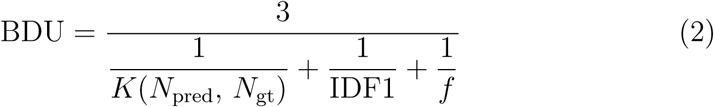

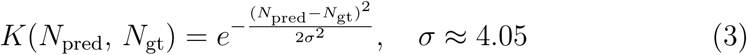

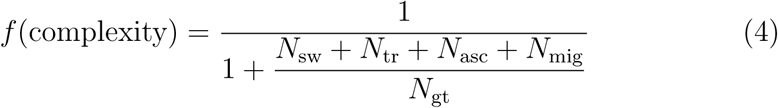

where *N*_pred_ is predicted specimens, *N*_gt_ is the ground-truth, *N*_sw_ is track switches, *N*_tr_ is transfers, *N*_asc_ is ascensions, and *N*_mig_ is migrations. Further detail on the count accuracy term *K*(*N*_pred_*, N*_gt_), and choice of *σ* in Appendix A. The BDU was developed independent of the JEPA-SORT architecture. An experiment justifying the BDU by correlating it with downstream biodensity estimate error can be found in Section 4.

### 2.4. Description of the Benchmark

The provided open-source benchmark contains twelve deep-sea ROV transects of benthic corals from Cabo Verde. The transects were recorded during a joint Instituto do Mar - OceanX mission in 2025. Ground-truth tracks in each transect were hand-labelled internally at OceanX.

### 2.5. Experimental Design

We benchmark four trackers: SORT, OC-SORT, DeepSORT, and DeepOC-SORT against JEPA-SORT on the twelve deep-sea ROV transects of benthic corals from Cabo Verde. DeepSORT and DeepOC-SORT are each tested with two embedding models, BioCLIP 2.5 (Gu et al., 2025) and DINOv3 (Siméoni et al., 2025). To assess performance across conditions, we augment the footage into eight configurations spanning a broad range of optical environments, each initializing an oracle detector from ground-truth with a unique combination of detection dropout rate, false-positive rate, stochastic false-positive propagation length, bounding-box area jitter, and effective frame rate. These parameters were sampled via Latin hypercube to construct an unbiased, representative set of conditions defined in Table 2. The alignment of each environment along false-positive clutter - detection stability axis can be found in Figure 1.

**Figure 1:**
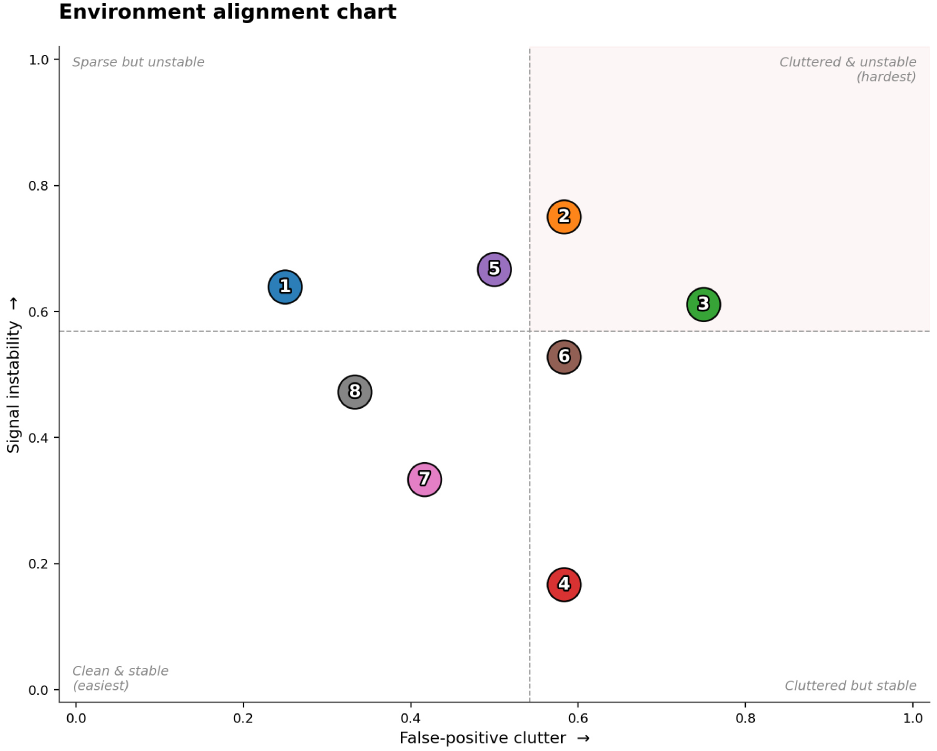
Environment alignment chart for the oracle-detector parameter settings.

**Table 2:**
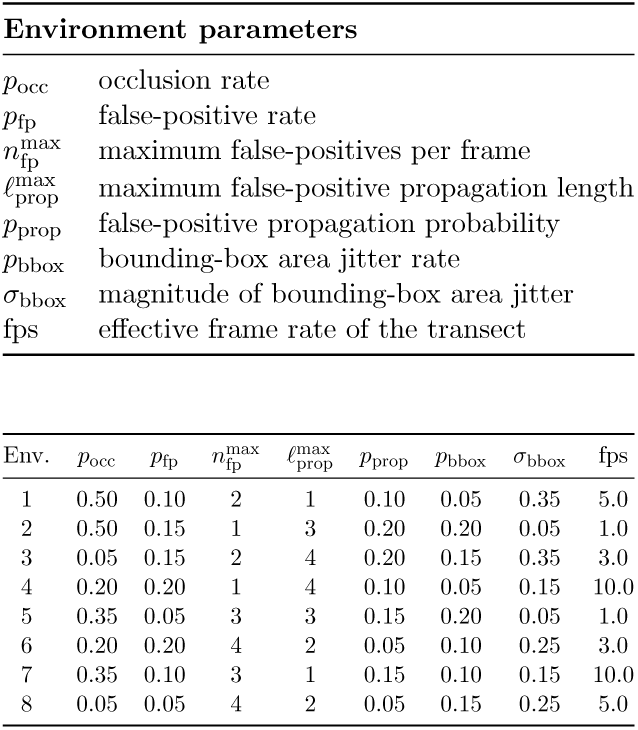
Oracle-detector environmental parameters.

| **Environment parameters** |  |  |  |  |  |  |  |
|---|---|---|---|---|---|---|---|
| p<sub>occ</sub> | occlusion rate |  |  |  |  |  |  |
| p<sub>fp</sub> | false-positive rate |  |  |  |  |  |  |
| n<sub>fp</sub><sup>max</sup> | maximum false-positives per frame |  |  |  |  |  |  |
| ℓ<sub>prop</sub><sup>max</sup> | maximum false-positive propagation length |  |  |  |  |  |  |
| p<sub>prop</sub> | false-positive propagation probability |  |  |  |  |  |  |
| p<sub>bbox</sub> | bounding-box area jitter rate |  |  |  |  |  |  |
| σ<sub>bbox</sub> | magnitude of bounding-box area jitter |  |  |  |  |  |  |
| fps | effective frame rate of the transect |  |  |  |  |  |  |
| Env. | p<sub>occ</sub> | p<sub>fp</sub> | n<sub>fp</sub><sup>max</sup> | ℓ<sub>prop</sub><sup>max</sup> | p<sub>prop</sub> | p<sub>bbox</sub> | σ<sub>bbox</sub> | fps |
| 1 | 0.50 | 0.10 | 2 | 1 | 0.10 | 0.05 | 0.35 | 5.0 |
| 2 | 0.50 | 0.15 | 1 | 3 | 0.20 | 0.20 | 0.05 | 1.0 |
| 3 | 0.05 | 0.15 | 2 | 4 | 0.20 | 0.15 | 0.35 | 3.0 |
| 4 | 0.20 | 0.20 | 1 | 4 | 0.10 | 0.05 | 0.15 | 10.0 |
| 5 | 0.35 | 0.05 | 3 | 3 | 0.15 | 0.20 | 0.05 | 1.0 |
| 6 | 0.20 | 0.20 | 4 | 2 | 0.05 | 0.10 | 0.25 | 3.0 |
| 7 | 0.35 | 0.10 | 3 | 1 | 0.15 | 0.10 | 0.15 | 10.0 |
| 8 | 0.05 | 0.05 | 4 | 2 | 0.05 | 0.15 | 0.25 | 5.0 |

The benchmark runs in two Monte Carlo stages, optimization and evaluation, in which each tracker is run repeatedly under stochastic oracle-detector perturbations. Optimization tunes hyperparameters for each MOT algorithm in each environment on a held-out training set of 2 transects, via a 200-trial random search; each trial scores a candidate configuration by the mean Biodensity Utility Metric over 10 runs (2 transects × 5 stochastic runs). The best configuration per tracker-environment pair is carried forward.

Evaluation applies these configurations to the test set of 10 remaining transects, running each tracker-environment-transect triplet five times. In both stages, random seeds vary across runs but are shared by run index, transect, and environment, ensuring reproducibility and identical perturbations across trackers. Scores are aggregated by arithmetic mean at the transect, environment, tracker, and overall levels, and reported in Section 3.

## 3. RESULTS

On the benchmark test split of ten ROV transects, JEPA-SORT outperforms SORT, OC-SORT, DeepSORT, and DeepOC-SORT on key metrics including IDF1, MOTA, and BDU. It also shows strong specimen-counting performance (0.784 ± 0.109, Figure 2), corresponding to a count error of ∼2.8 specimens. On count accuracy alone it is narrowly edged out by both DeepOC-SORT variants (0.805±0.066 and 0.800±0.08, ∼2.7 specimens), but JEPA-SORT remains the strongest tracker overall on both traditional metrics and BDU. A paired-difference plot, Figure 3, confirms JEPA-SORT’s lead on all three headline metrics; pairing within environment-transect-seed blocks removes between-transect and seed variance, tightening the confidence intervals.

**Figure 2:**
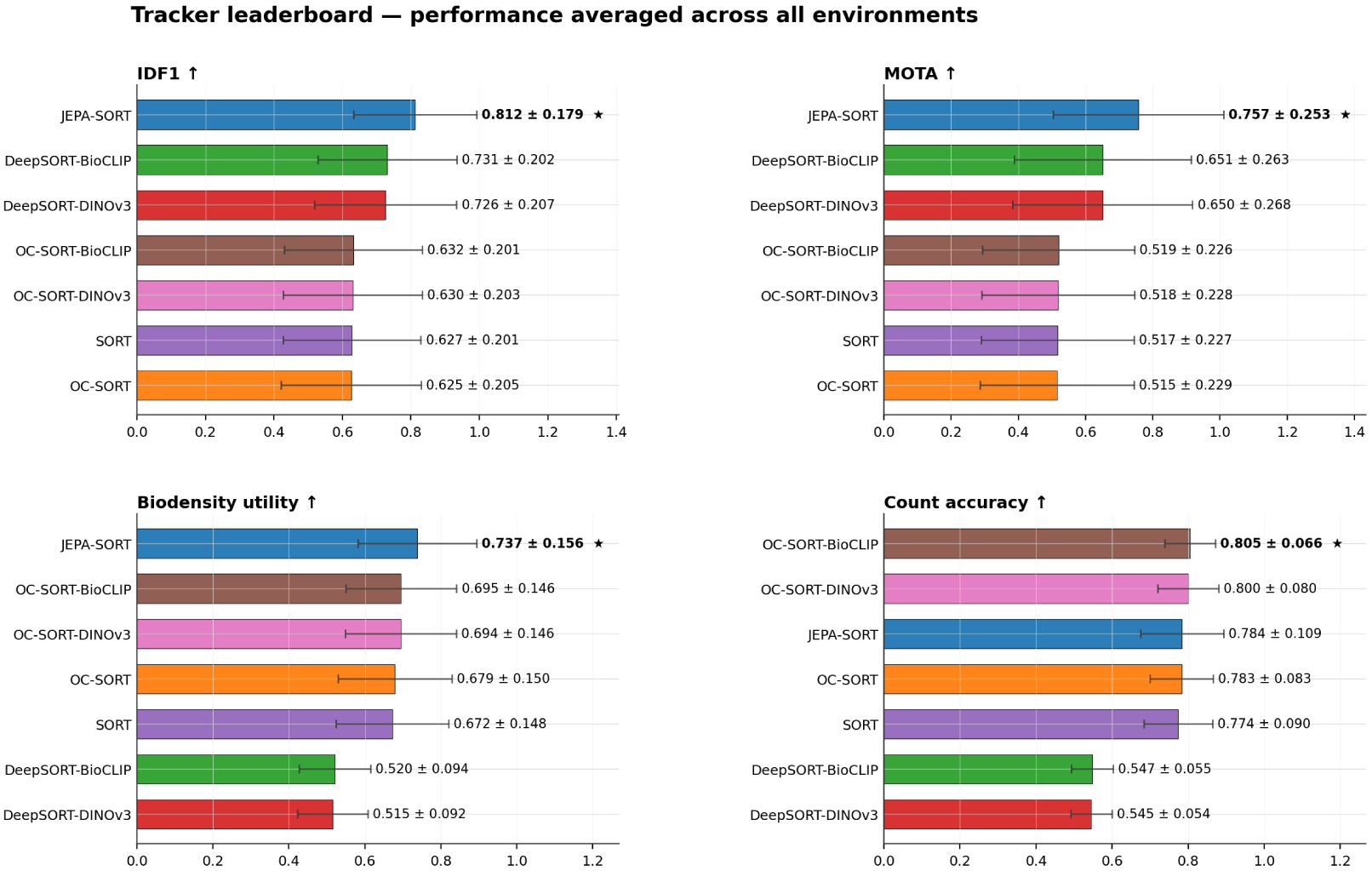
Tracker performance on key MOT metrics averaged across environments and transects.

**Figure 3:**
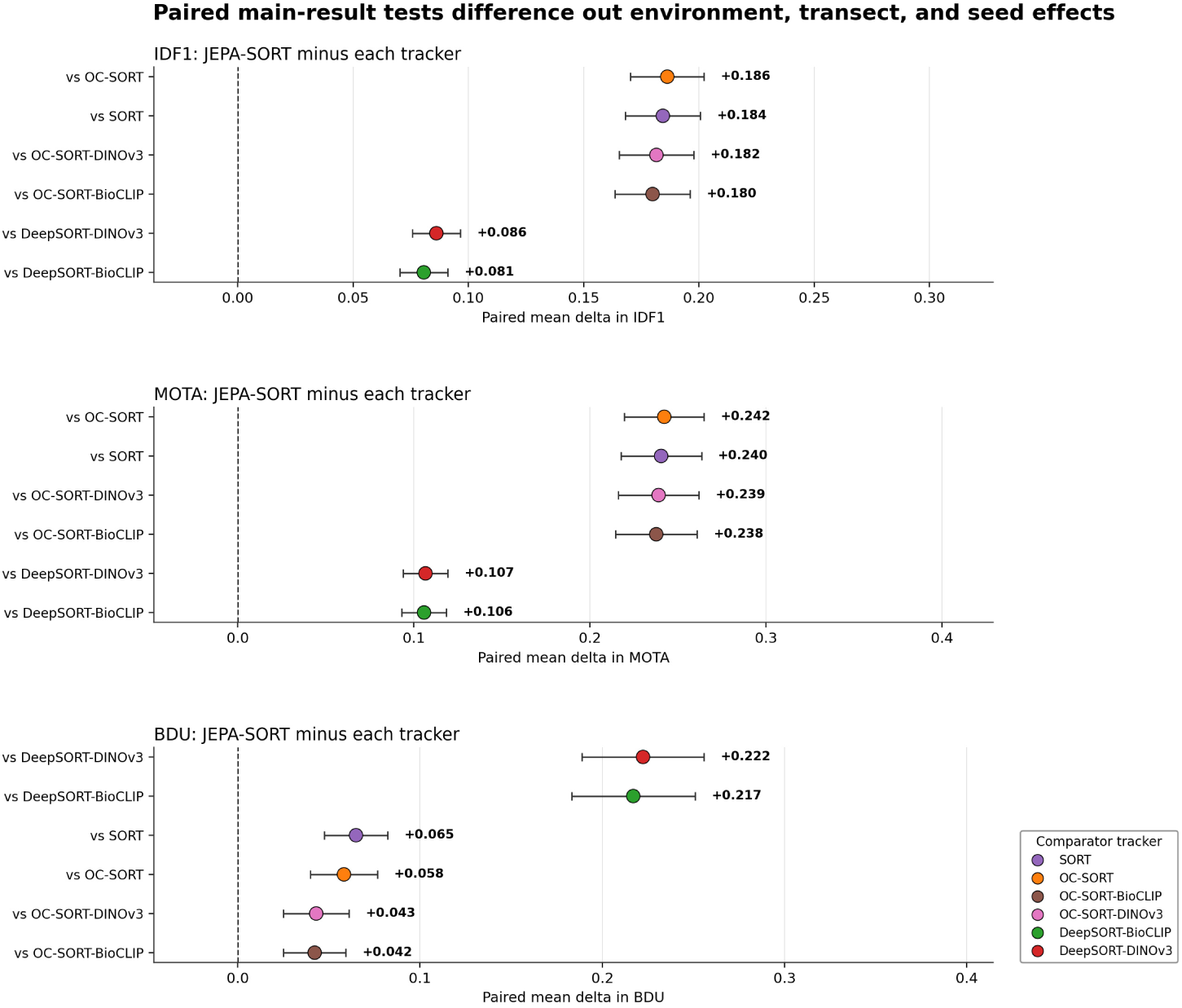
Paired difference of JEPA-SORT vs other benchmark trackers averaged across (environment, transect, run) tuples.

Performance among SORT, OC-SORT, DeepSORT, and DeepOC-SORT variants is more mixed. DeepSORT outperforms SORT, OC-SORT, and DeepOC-SORT on IDF1 and MOTA. However, the trend is reversed on BDU and raw count accuracy, with the DeepSORT variants being the least performant across the entire set of trackers. This highlights how performance on traditional metrics isn’t necessarily aligned with performance in the context of downstream ecological modeling.

As shown in Figure 4, at the environment-transect level, JEPA-SORT retains its overall advantage, revealing the tracker’s relative strengths while also exposing unique failure modes absent in the other trackers. Notably, although it leads on IDF1, BDU, and MOTA, it underperforms on the lower-level scores that compose them: ID switches, track transfers, track ascensions, and track migrations (see Figure 5). DeepOC-SORT leads on ID switches and ascensions, DeepSORT leads on transfers, and SORT leads on migrations. However, this apparent paradox resolves once recall is taken into account.

**Figure 4:**
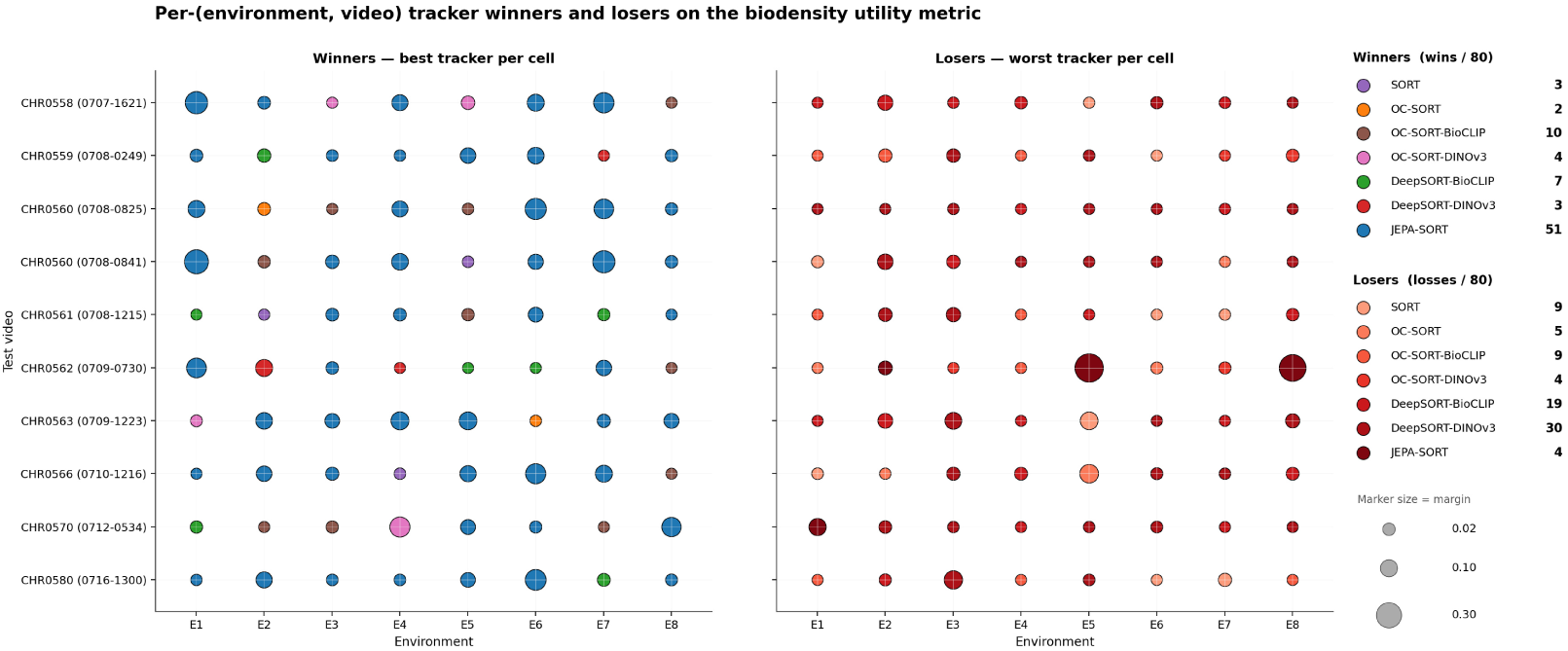
Winner and loser performance margin across environment-transect pairs.

**Figure 5:**
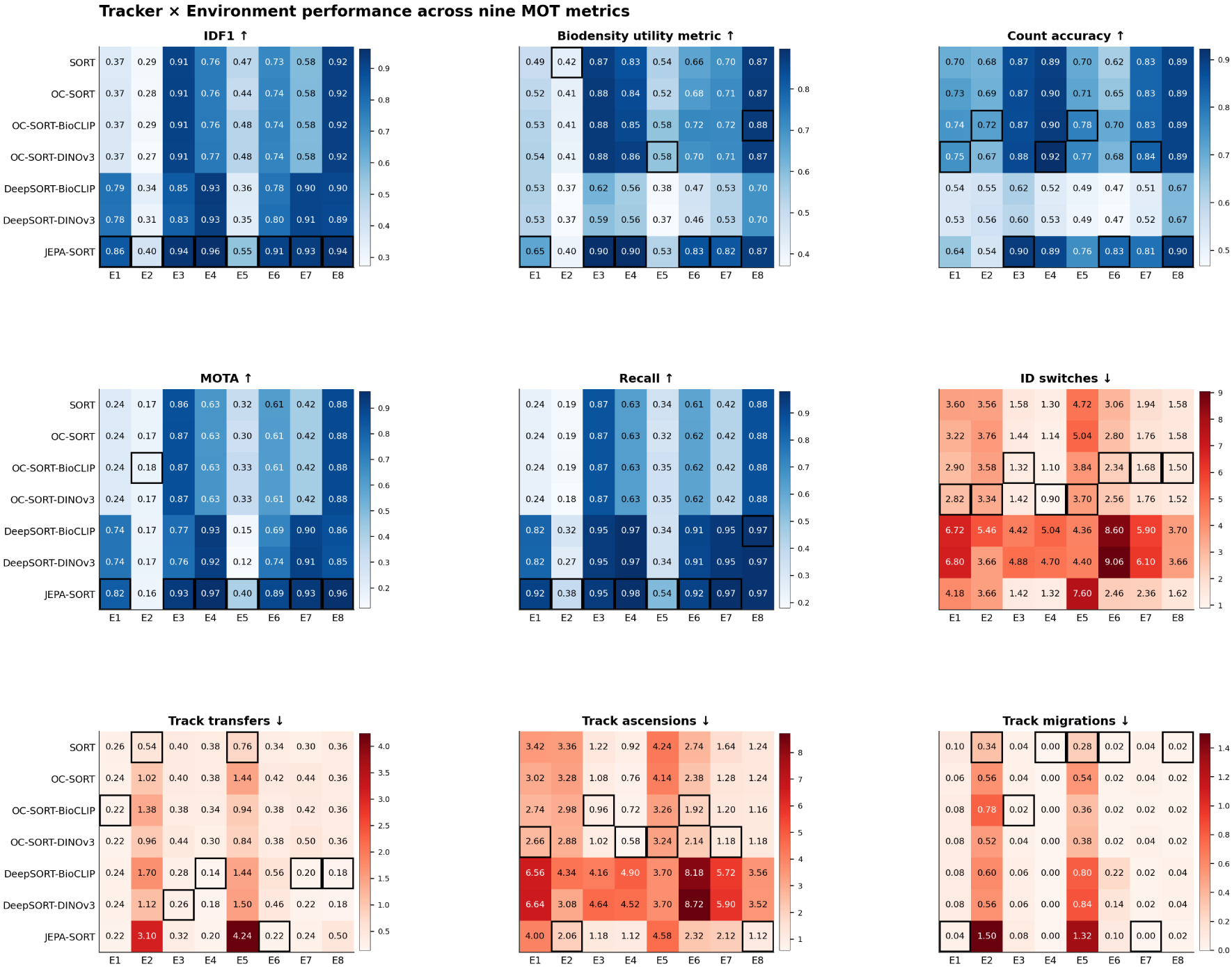
Transect-averaged tracker performance on MOT metrics across environments.

To normalize by recall, we divide a given tracker-environment count by its associated recall rate. For example, to normalize ID switches by recall, we simply divide the components of the ID switches matrix by their corresponding recall values.

As shown in Figure 6, when normalized by recall JEPA-SORT outperforms on switches and ascensions, and is competitive on transfers and migrations. In other words, per unit recall, JEPA-SORT actually maintains association better than, or equal to SORT, OC-SORT, DeepSORT and DeepOC-SORT. For example, while DeepOC-SORT leads on raw ID switches, JEPA-SORT is generally the most performant on recall-normalized ID switches. JEPA-SORT’s apparent underperformance on raw track-quality scores is therefore an artifact of its higher recall: by maintaining tracks over longer durations, it also exposes more opportunities for switches, transfers, ascensions, and migrations to occur.

**Figure 6:**
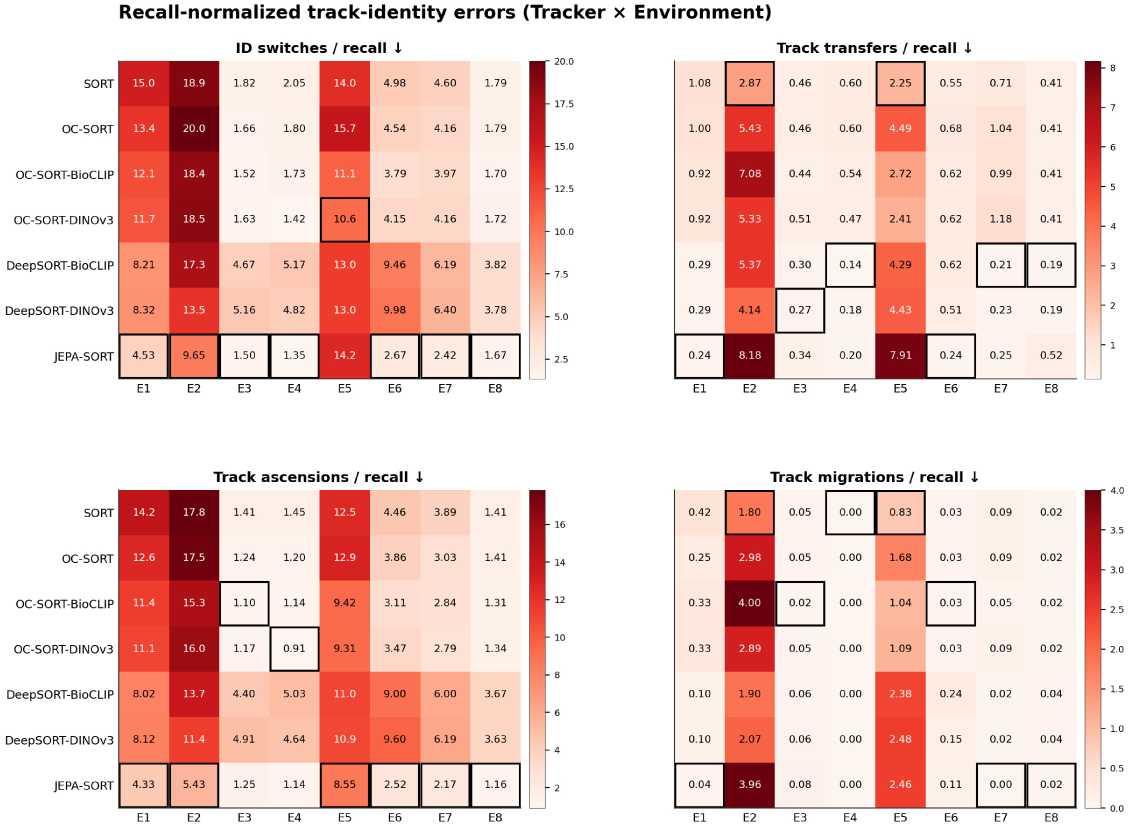
Transect-averaged tracker performance on MOT metrics across environments, normalized by recall.

Despite these strengths, the data also reveals a failure mode unique to JEPA-SORT. Per Figure 4, while it achieves the highest BDU score on ∼64% of the eighty transect-environment pairs, its failures are concentrated and severe: on pairs CHR0562 (0709-0730)-E5 and -E8, its BDU falls more than 0.30 below the second-worst tracker.

Notably, CHR0562 (0709-0730) is essentially still, unlike the training (optimization) split, which exhibits significant ego-motion. Given JEPA-SORT’s relatively large number of hyperparameters, the tracker may have overfit to the training data and failed on this out-of-distribution transect. Why it failed specifically on -E5 and -E8, while remaining in-family with the other trackers on -E1 through -E4, -E6, and -E7, we leave to future work. The following ablations instead focus on separating the contributions of V-JEPA 2.1 from the broader JEPA-SORT architecture and show that video world-model embeddings help discriminate between visually homogeneous specimens in deep-sea environments.

### 3.1. Ablations

As the JEPA-SORT algorithm demonstrates robust performance on the 12-transect benchmark of deep-sea corals, we conduct the following two ablations to identify the underlying driver of its improved associative ability.

#### 3.1.1. Relative contribution of JEPA-SORT architecture and V-JEPA 2.1

To isolate the relative contribution of the JEPA-SORT architecture and the V-JEPA 2.1 embeddings to the overall tracker performance, we substituted DINOv3 for V-JEPA 2.1 within the JEPA-SORT architecture’s associative step, and re-evaluated on the same set of transect-environment pairs. The results show the architecture and embedding components play distinct and complementary roles, although the decisive performance gains are driven by the usage of V-JEPA 2.1’s video world-model embeddings.

Holding the embedding fixed at DINOv3, architecture choice moves a tracker along a recall/count-accuracy trade-off rather than improving both jointly. As shown in Figure 7B DeepSORT’s architecture maximizes recall at the cost of count accuracy, OC-SORT’s architecture protects count accuracy at the cost of recall, and JEPA-SORT’s architecture lands between the two on both axes. Once the embedding model is held constant, JEPA-SORT does not outperform OC-SORT on BDU.

**Figure 7:**
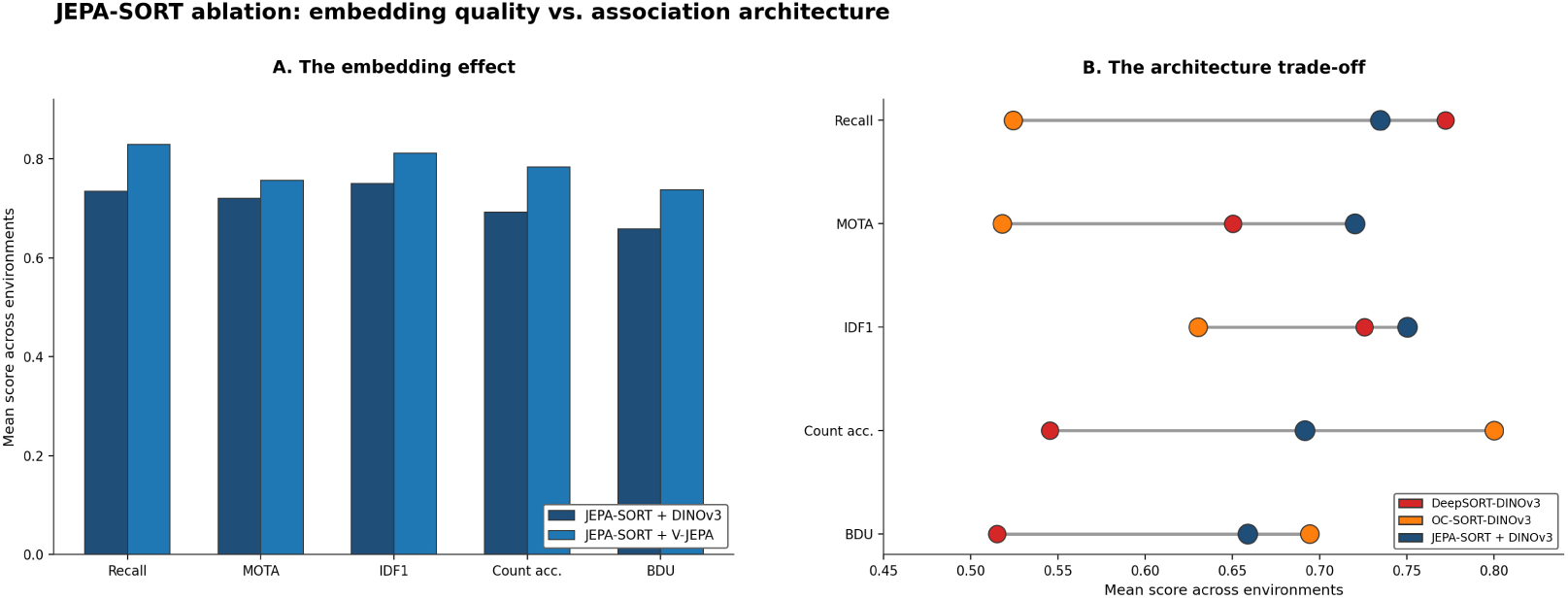
A: Relative gain of V-JEPA 2.1 vs DINOv3 in the JEPA-SORT architecture, B: Relative architecture performance using DINOv3.

Figure 7A tells a different story. Holding the architecture fixed at JEPA-SORT, using V-JEPA 2.1 as the embedding model vs DINOv3 lifts recall, MOTA, IDF1, count accuracy, and BDU simultaneously, with no corresponding trade-off. V-JEPA 2.1 raises JEPA-SORT’s balanced count accuracy and recall metrics together. Practically, the JEPA-SORT algorithm should therefore be understood as a harness necessary to utilize video world-model embeddings, with decisive performance gains driven by the discriminative power of those embeddings.

#### 3.1.2. Embedding model identity accuracy

To better understand the discriminative strength of V-JEPA 2.1 relative to DINOv3 and BioCLIP 2.5, we calculate each model’s nearest-neighbor identity accuracy. In a given transect, for each ground-truth detection *d_i_* we find the detection *d_j_* with the greatest cosine similarity and record whether *d_j_* belongs to the same ground-truth track. To prevent long tracks from dominating, we compute the NN accuracy within each track, then average these per-track accuracies across the transect, yielding a track-balanced accuracy. This is more informative than comparing average intra- and inter-track cosine similarity: because V-JEPA 2.1, BioCLIP 2.5, and DINOv3 project onto embedding spaces with different inherent scales of similarity, raw similarities are not comparable across models.

As illustrated in Figure 8 V-JEPA 2.1 leads on track-balanced nearest neighbor identity accuracy in 66.7% of transects, with DINOv3 at 16.7%, and BioCLIP at 0.0%. The remaining 17% of transects are ties. Notably, V-JEPA 2.1’s lead is most significant on transects with high specimen visual homogeneity such as CHR0561 (0708-1215), CHR0580 (0716-1300), CHR0560 (0708-0825), and CHR0570 (0712-0534), where per-frame appearance features are insufficient to distinguish between unique specimens and spatiotemporal embeddings offer an associative edge. Frames from these transects are shown in Figure 9.

**Figure 8:**
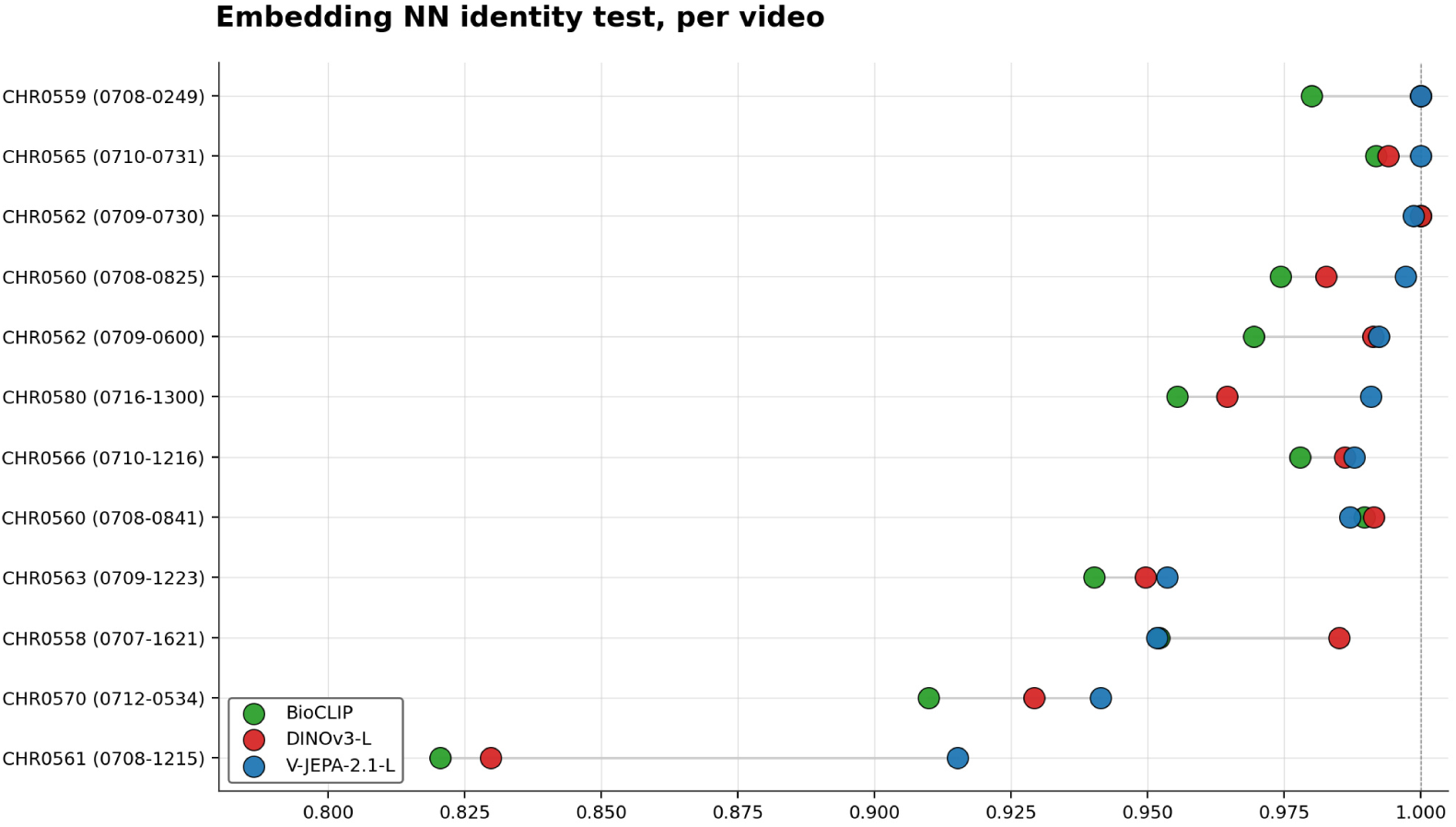
Track-balanced nearest-neighbor (NN) accuracy per transect.

**Figure 9:**
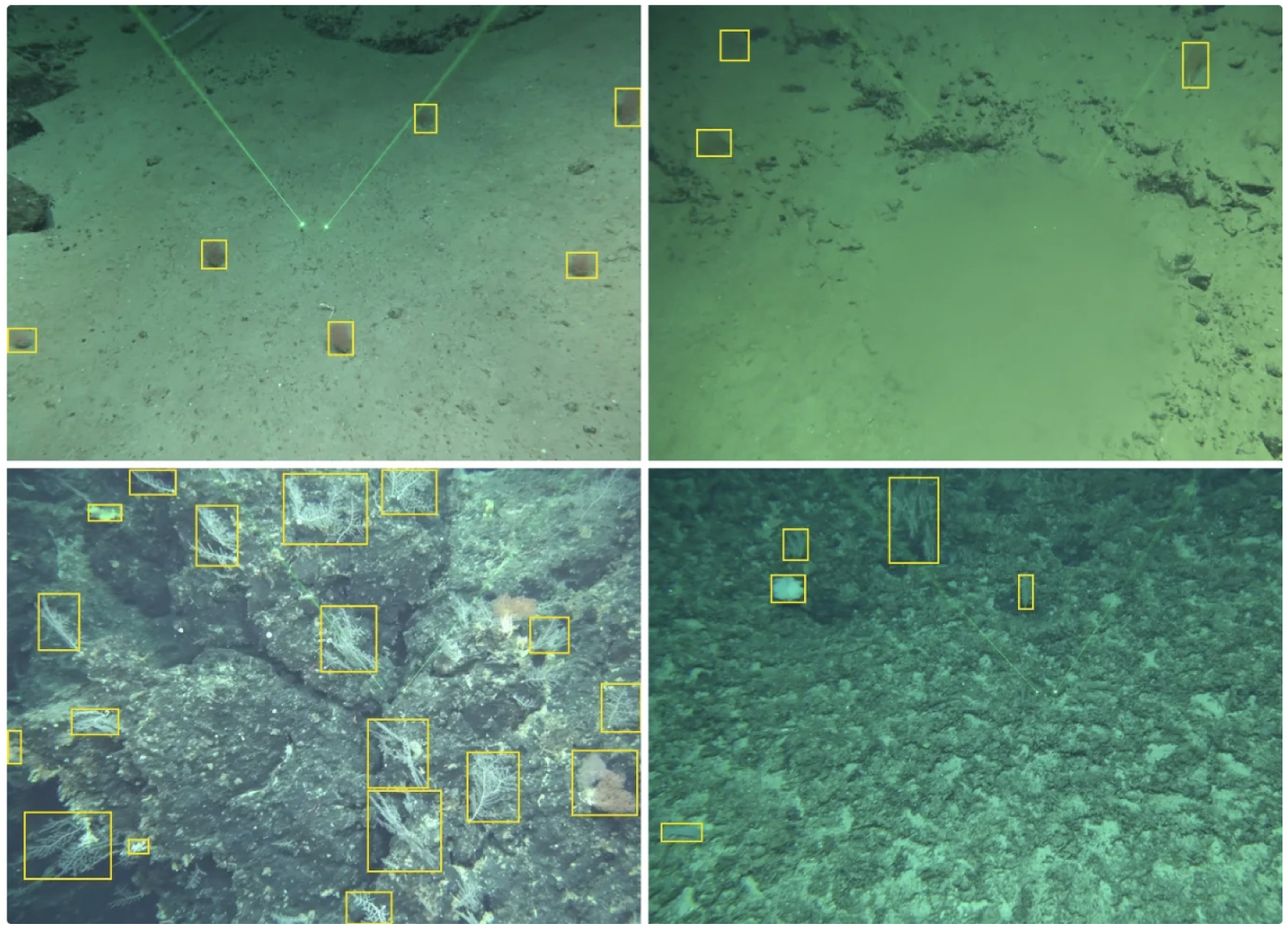
Clockwise: Sample frame from CHR0561 (0708-1215), CHR0580 (0716-1300), CHR0560 (0708-0825), and CHR0570 (0712-0534)

In comparison, the transects where DINOv3 leads such as CHR0558 (0707-1621) typically have relatively low specimen visual homogeneity, where appearance alone is sufficient to establish identity (see Figure 10).

**Figure 10:**
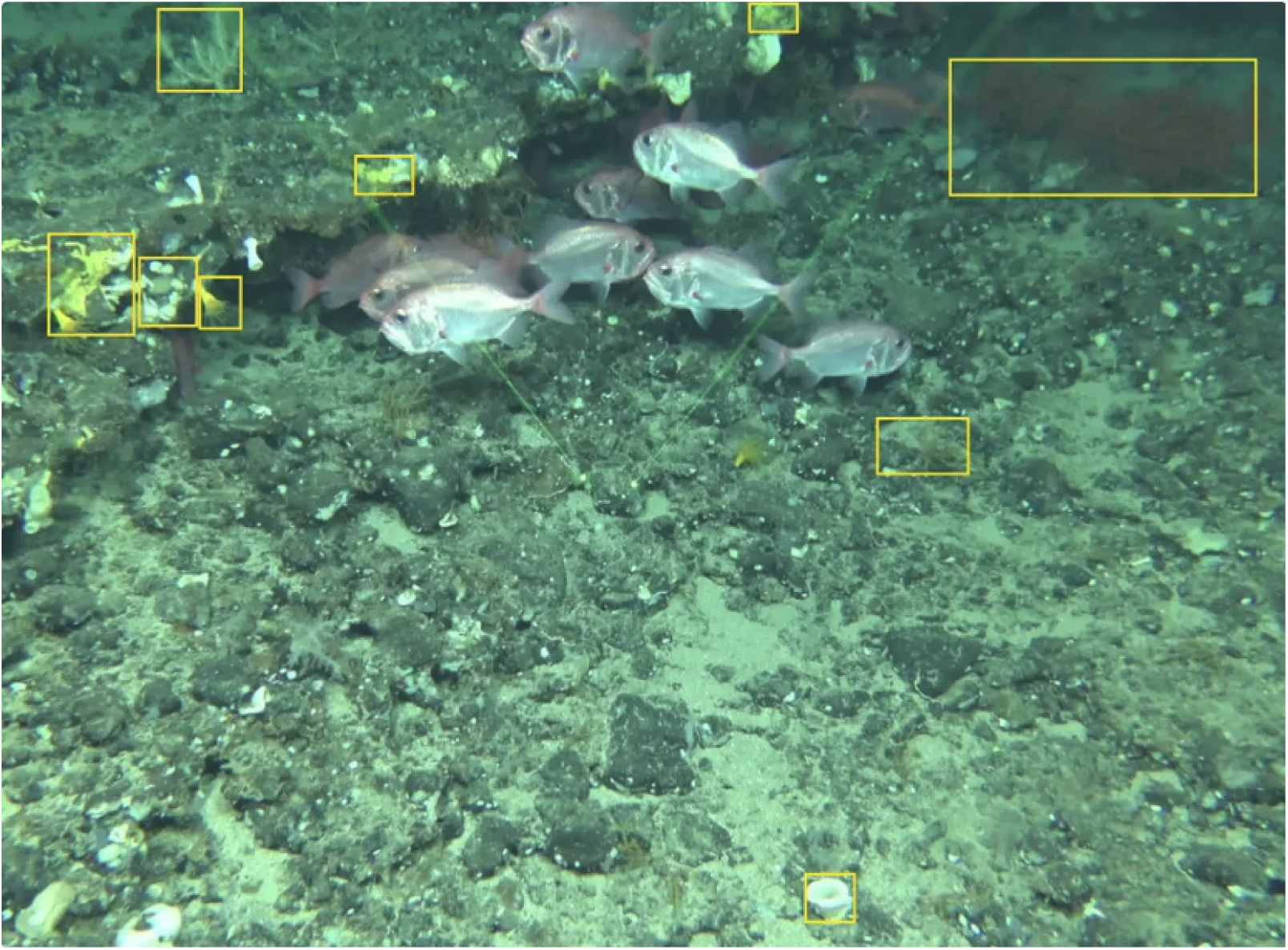
Sample frame from CHR0558 (0707-1621)

These results show that the key advantage of V-JEPA 2.1’s spatiotemporal embeddings is that they offer an associative edge on the most visually homogeneous footage. Deep association alone, without addressing the appearance-discrimination bottleneck that plagues visually homogeneous deep-sea footage, does not improve downstream biodensity estimation.

## 4. ANALYSIS OF THE BDU

To validate BDU against a real ecological analysis, we used each tran-sect’s paired laser scalers (spaced 10 cm apart) to calculate an area density estimate (specimens per unit area) and measured each tracker’s density error as absolute relative error between predicted and ground-truth area density, 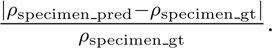 The area estimation process is illustrated in Figure 11.

**Figure 11:**
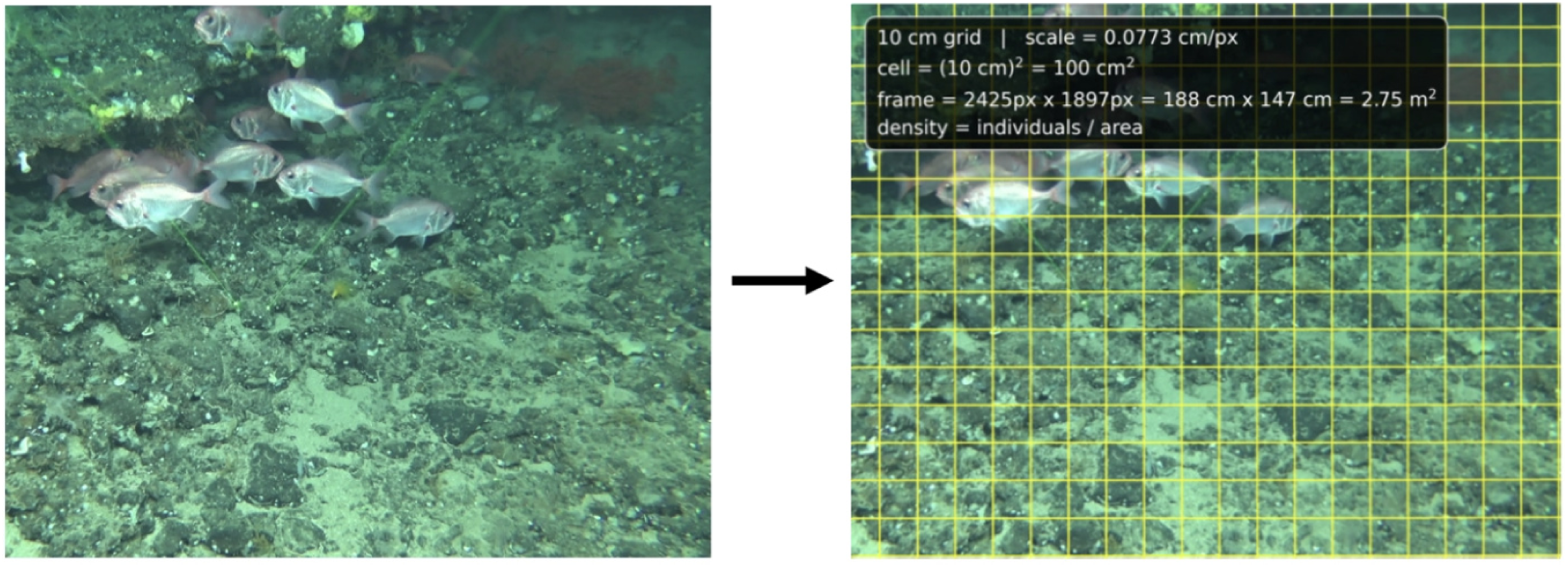
Illustration of area estimation based on 10 cm laser spacers.

As shown in Figure 12, BDU’s ranking of trackers correlates strongly with their relative areal-density error (mean |*ρ*| = 0.72), substantially exceeding the standard identity metrics IDF1 and MOTA (mean |*ρ*| = 0.56 and 0.56). This is both desirable and to be expected, since areal density is a function of raw specimen count error which is a contributing term in eq (3) and by extension the BDU. However, for ecological applications, track consistency is of comparable importance to raw count accuracy, as a sustained sequence of detections per individual yields multiple views which increases the opportunity for identification.

**Figure 12:**
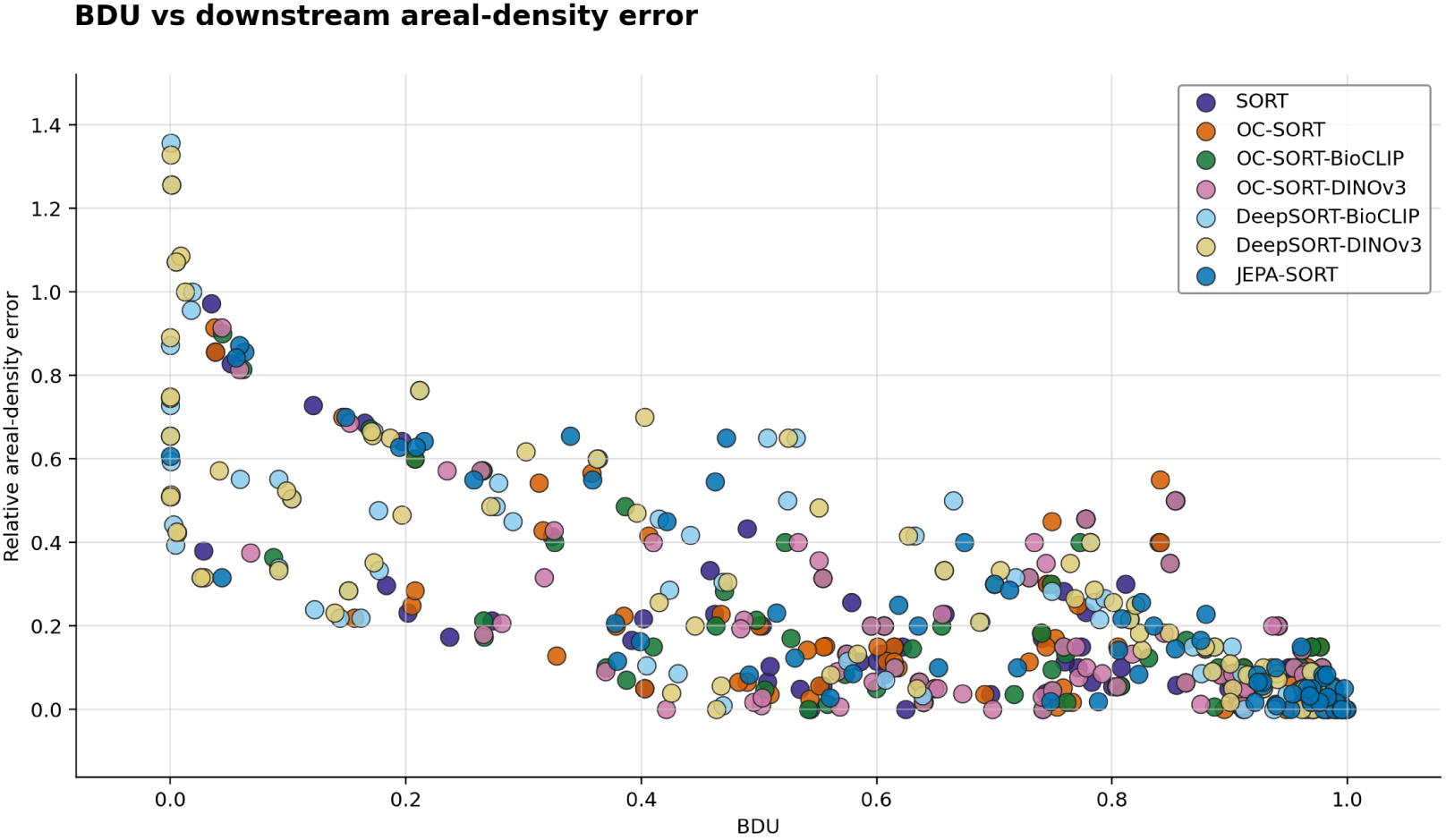
Scatter plot of area-density error versus BDU across transect-environment pairs.

To assess this second focus, we correlated each metric against track-coverage outcomes at the pooled tracker-environment level (Table 3), and the pattern is complementary to the density result. Count accuracy, the best predictor of density error, is nearly blind to coverage (*r* = −0.007 with mostly-tracked individuals), whereas IDF1 and MOTA - weak on density – are the strongest coverage predictors (*r* = 0.828 and 0.870). No single standard metric captures both axes. BDU reflects each by construction: it predicts density error far better than IDF1 and MOTA, and is positively associated with coverage (*r* = 0.41) where count accuracy is not. It leads on neither axis but is the only metric that meaningfully reflects both - the balance an ecological metric requires, since abundance and consistent identity are both essential downstream.

**Table 3:**
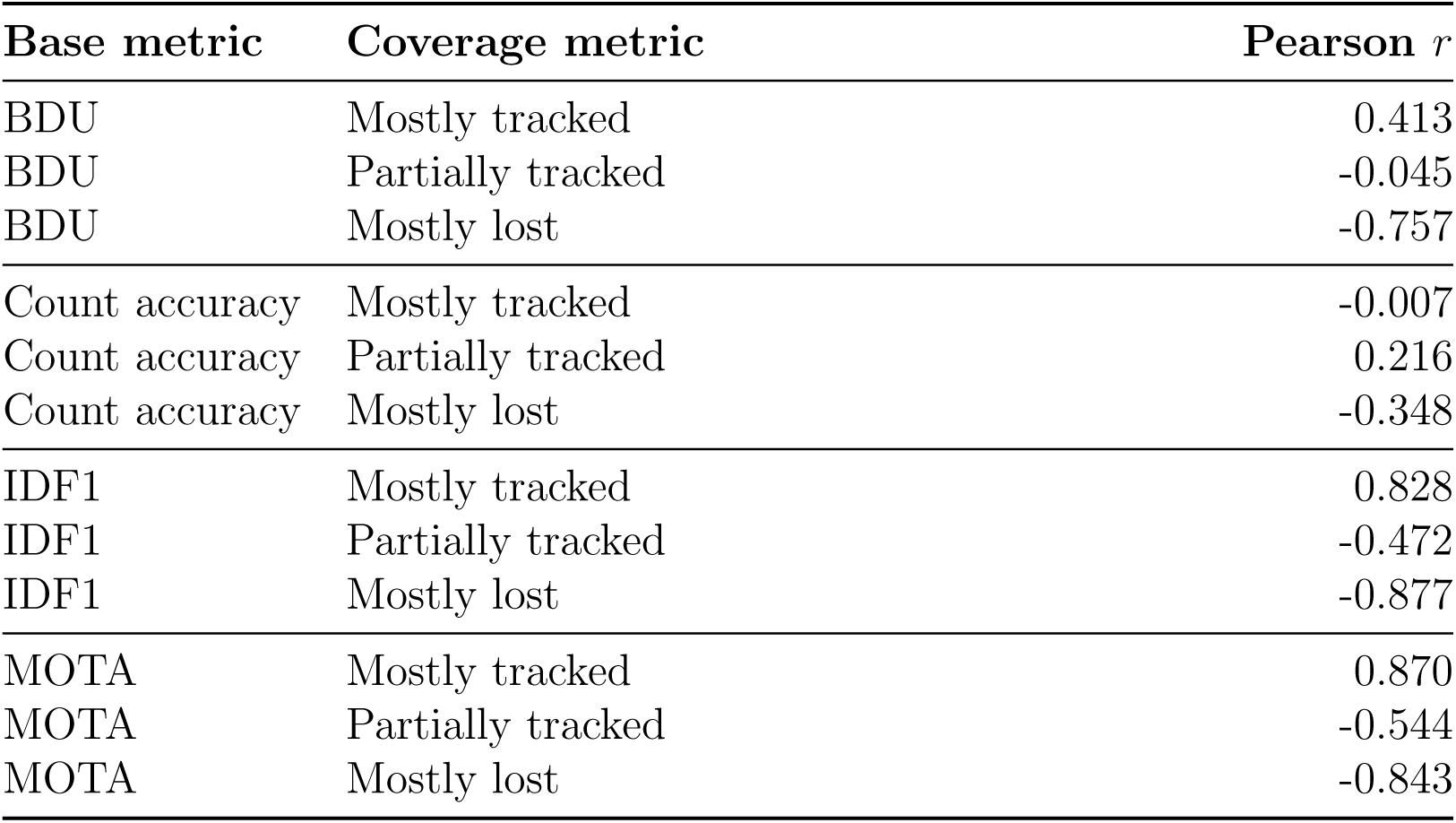
Pearson correlations between base tracking metrics and coverage metrics.

| **Base metric** | **Coverage metric** | **Pearson *r*** |
|---|---|---|
| BDU | Mostly tracked | 0.413 |
| BDU | Partially tracked | -0.045 |
| BDU | Mostly lost | -0.757 |
| Count accuracy | Mostly tracked | -0.007 |
| Count accuracy | Partially tracked | 0.216 |
| Count accuracy | Mostly lost | -0.348 |
| IDF1 | Mostly tracked | 0.828 |
| IDF1 | Partially tracked | -0.472 |
| IDF1 | Mostly lost | -0.877 |
| MOTA | Mostly tracked | 0.870 |
| MOTA | Partially tracked | -0.544 |
| MOTA | Mostly lost | -0.843 |

## 5. DISCUSSION

Ablations indicate that the primary bottleneck in deep-sea benthic footage is appearance discrimination rather than kinematics, and that spatiotemporal appearance embeddings from a video world-model can help overcome it: by encoding a specimen’s trajectory across frames alongside its physical appearance, they offer an associative edge where per-frame features are too weak to separate visually homogeneous specimens. Holding the embedding fixed, architecture choice moves a tracker along a recall/count-accuracy trade-off, whereas substituting V-JEPA 2.1 for DINOv3 lifts recall, MOTA, IDF1, count accuracy, and BDU simultaneously, with the largest gains on the most visually homogeneous transects. Practically, deep association alone is insufficient in this environment; what matters is the discriminative power of the embedding on specimens that are difficult to tell apart. Notably, DeepSORT is strong on IDF1 and MOTA while being the least performant on BDU and count accuracy, which highlights how performance on traditional metrics isn’t necessarily aligned with performance in the context of downstream ecological modeling.

JEPA-SORT belongs to the SORT family of tracking-by-detection algorithms. The original SORT (Bewley et al., 2016) performs purely kinematic association via a Kalman filter and IoU matching. DeepSORT extended it by integrating appearance embeddings for detection re-identification (Wojke et al., 2017), and OC-SORT introduced observation-centric momentum, which penalizes implausible direction changes and re-associates tracks against actual observations rather than accumulated filter predictions to recover identities through detection dropout (Cao et al., 2023). DeepOC-SORT combines both directions, pairing OC-SORT’s observation-centric association with appearance embedding (Maggiolino et al., 2023). Across this family, association relies on kinematics, per-frame appearance embeddings, or both - which our ablations indicate is why the family plateaus on visually homogeneous footage; we instead draw on the Joint-Embedding Predictive Architecture line of work, originally proposed by (LeCun, 2022) and realized for video in V-JEPA 2 and V-JEPA 2.1, which to our knowledge has not previously been used for MOT association.

Within the underwater domain, tracking research has largely focused on pelagic fish in aquaculture or shallow-reef settings. FishMOT (Liu et al., 2024) uses a temporal re-find module to recover identities through detection gaps under turbidity, while a recent transformer-based tracker handles trajectory-aware deep association implicitly inside the network, rather than injecting pretrained video embeddings into an explicit SORT-style association step as we do (Li and Han, 2026). Earlier underwater vision benchmarks such as the Brackish dataset established detection under varying visibility as a core challenge, but targeted fish and marine animals in shallow water rather than the sessile benthic organisms and tracking task we address (Pedersen et al., 2019). The most similar benchmark, DeepSeaMOT, provides only two benthic transects and does not compare trackers (Barnard et al., 2025).

### 5.1. Scope and Future Work

Several aspects of this study define the scope of its conclusions and suggest natural extensions. The oracle detector was chosen to isolate tracker behavior from detector-specific error, giving controlled and reproducible variation across optical environments; pairing JEPA-SORT with a trained detector is the natural next step toward deployment. The benchmark likewise covers benthic corals in a single region, and extending it across regions, taxa, and density regimes would establish how broadly the advantage of world-model embeddings holds. BDU’s count tolerance and term weighting were calibrated for this setting and can be adapted where an application weights abundance and identity consistency differently. Finally, because the specimens here are sessile, the 16-frame window encodes viewpoint variation across ego-motion. Future work could include the analysis of ROV transects showing non-sessile organisms.

## 6. CONCLUSION

This paper provides a benchmark of twelve deep-sea ROV transects surveying benthic corals, evaluates four standard MOT algorithms across eight simulated optical environments, and introduces JEPA-SORT, a tracker that injects V-JEPA 2.1 spatiotemporal embeddings into SORT-style cascading association. We also define the Biodensity Utility Metric (BDU), a composite that ties tracker performance to the requirements of downstream ecological modeling.

JEPA-SORT is the strongest tracker in the benchmark, leading on IDF1, MOTA, and BDU while remaining competitive on specimen count, and is the only tracker to maintain both high recall and stable identity association across optical environments - its higher raw track-quality error counts prove, once normalized by recall, to be an artifact of sustaining tracks longer rather than of weaker association. Ablations indicate this advantage stems primarily from the discriminative power of world-model embeddings on visually homogeneous footage. No single standard metric captures both axes of ecological utility: count accuracy predicts areal-density error but is blind to track coverage, while IDF1 and MOTA predict coverage but not density; BDU is the only metric that meaningfully reflects both. JEPA-SORT’s clearest failure mode - apparent overfitting on a small optimization split - points to a natural next step: expanding the benchmark to support a larger training set.

## APPENDIX A

### Count accuracy term K(N_pred_, N_gt_)

The count-accuracy term, *K*(*N_pred_, N_gt_*), is an RBF kernel, chosen for interpretability and invariance to relative error: it is symmetric about zero, and unlike raw count difference, a given error *N_pred_*_−_*_gt_* produces the same score regardless of *N_gt_*. This is easily interpretable at the transect level, but the kernel’s Gaussian curvature complicates aggregation across environments. By Jensen’s inequality, *f* (*E*[*X*]) ≥ *E*[*f* (*X*)] for concave *f*, so the count accuracy of the mean error, *K*(*E*[*N_pred_* − *N_gt_*]), may exceed the mean count accuracy, *E*[*K*(*N_pred_* − *N_gt_*)]. We adopt the latter: count accuracy is computed per tracker-environment-transect triplet and averaged across environments, i.e. we report *E*[*K*(*N_pred_* − *N_gt_*)].

### Choice of σ

The value *σ* ≈ 4.05 was set empirically to produce a graceful penalty curve over the range of count errors observed in our transects. Under this choice, a miscount of one organism retains 97 % accuracy, two retains 89 %, and accuracy falls to 61 % at four and 14 % at eight - penalizing small errors lightly while degrading smoothly for larger deviations.

